# Rapid evolution of complete dosage compensation in *Poecilia*

**DOI:** 10.1101/2021.02.12.431036

**Authors:** David C.H. Metzger, Benjamin A. Sandkam, Iulia Darolti, Judith E. Mank

## Abstract

Dosage compensation balances gene expression between the sexes in systems with diverged heterogametic sex chromosomes. Theory predicts that dosage compensation should rapidly evolve in parallel with the divergence of sex chromosomes to prevent the deleterious effects of dosage imbalances that occur as a result of sex chromosome divergence. Examples of complete dosage compensation, where gene expression of the entire sex chromosome is compensated, are rare and have only been found in relatively ancient sex chromosome systems. Consequently, very little is known about the evolutionary dynamics of complete dosage compensation systems. We recently found the first example of complete dosage compensation in a fish, *Poecilia picta*. We also found that the Y chromosome degraded substantially in the common ancestor of *P. picta* and their close relative *Poecilia parae*. In this study we find that *P. parae* also have complete dosage compensation, thus complete dosage compensation likely evolved in the short (∼3.7 my) interval after the split of the ancestor of these two species from *P. reticulata*, but before they diverged from each other. These data suggest that novel dosage compensation mechanisms can evolve rapidly, thus supporting the longstanding theoretical prediction that such mechanisms arise in parallel with rapidly diverging sex chromosomes.

**SIGNIFICANCE STATEMENT:** In species with XY sex chromosomes, females (XX) have as many copies of X-linked genes compared to males (XY), leading to unbalanced expression between the sexes. Theory predicts that dosage compensation mechanisms should evolve rapidly as X and Y chromosomes diverge, but examples of complete dosage compensation in recently diverged sex chromosomes are scarce, making this theory difficult to test. Across Poeciliid species the X and Y chromosomes have recently diversified. Here we find complete dosage compensation evolved rapidly as the X and Y diverged in the common ancestor of *Poecilia parae* and *P. picta*, supporting that novel dosage compensation mechanisms can evolve rapidly in tandem with diverging sex chromosomes. These data confirm longstanding theoretical predictions of sex chromosome evolution.

## INTRODUCTION

In organisms with heterogametic sex determination, the Y chromosome diverges from the X when recombination between them is suppressed (Furman et al. 2020). The same process holds for the Z and W chromosomes, but we focus here on male heterogametic systems. Degradation of the Y chromosome can lead to pseudogenization and gene loss resulting in females (XX) having twice as many copies of genes on the sex chromosome compared to males (XY). Because genes are normally expressed equally from both copies of a chromosome, males would only have half the expression of X-linked loci (Ohno 1967; Gu & Walters 2017), leading to a dosage imbalance with expression of genes on the autosomes. To resolve this issue, many organisms have evolved mechanisms to equalize expression levels of these sex chromosome genes, known as dosage compensation (Ohno 1967). Dosage compensation mechanisms are thought to evolve rapidly in parallel with Y degradation (Ohno 1967), however, the majority of sex chromosomes with dosage compensation are relatively old making it difficult to determine if dosage compensation can evolve in rapidly diverging sex chromosome systems.

Dosage compensation can either act by modifying expression on a gene-by-gene basis (incomplete dosage compensation) or by modifying expression along the entire chromosome (complete dosage compensation). Complete dosage compensation is predicted to arise for sex chromosomes that are rapidly diverging and experiencing extensive gene loss or pseudogenization, and has been more commonly found in male-heterogametic systems (XY) (Mullon et al. 2015; Wilson Sayres & Makova 2011). The most well characterized example for the rapid evolution of complete dosage compensation is in *Drosophila* where complete dosage compensation followed the emergence and divergence of a new XY sex chromosome system (Marín et al. 1996). The emergence of dosage compensation on neo-sex chromosomes in *Drosophila* is the result of evolution coopting extant dosage compensation mechanisms that predate the origin of the *Drosophila* genus (Marín et al. 1996). While dosage compensation can clearly evolve rapidly, it is unknown if complete dosage compensation can evolve rapidly when it is not present in close relatives.

Fish exhibit a high rate of sex chromosome turnover, and although there are some species with incomplete dosage compensation (eg. sticklebacks, flatfish, and rainbow trout) (White et al. 2015; Shao et al. 2014; Hale et al. 2018) complete dosage compensation appears to be rare. We recently identified the first example of complete dosage compensation in a fish; *Poecila picta. P. picta* is a close relative to the guppy (*Poecila reticulata*) (Darolti et al. 2019) that shares the same XY system that originated 18.48-26.08 Mya (Darolti et al. 2019; Rabosky et al. 2018). In *P. reticulata*, the X and Y have remained largely homomorphic, with little evidence of gene loss on the Y, and no need for dosage compensation (Darolti et al. 2019). However, since their split ∼18.4 Mya (Rabosky et al. 2018) the *P. picta* Y has diverged substantially from the X across nearly the entire chromosome and evolved complete dosage compensation (Darolti et al. 2019).

Here we take a comparative approach to narrow the timing of the evolution of complete dosage compensation by testing for dosage compensation in *P. parae* a sister taxon to *P. picta*. We recently characterized the sex chromosomes of *P. parae*, including five discrete Y haplotypes that control the five male morphs of this species (Sandkam et al. 2020). Importantly we found XY divergence across all five *P. parae* Ys was the same as XY divergence in *P. picta*, indicating the Y diverged from the X in the ∼3.7 my interval spanning the split of the *P. picta – P. parae* from the common ancestor with *P. reticulata* ∼18.4 mya, and prior to the split of *P. picta* and *P. parae* from each other ∼14.7 mya (Rabosky et al. 2018). Therefore, if *P. parae* also has complete dosage compensation, then dosage compensation evolved rapidly after XY chromosome divergence over a period of less than 3.7 million years (Figure 1).

**Figure 1.**
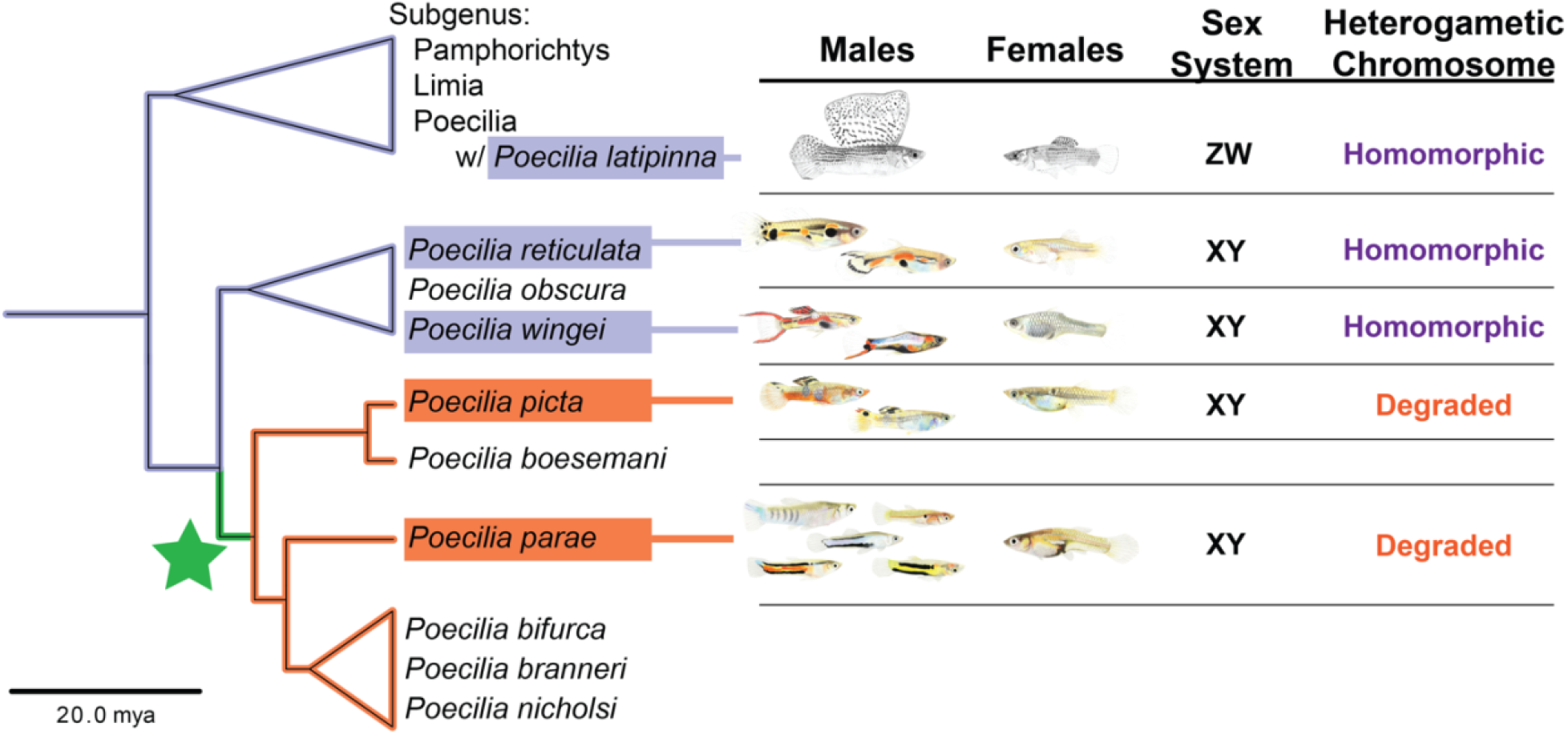
A phylogeny of Poecilia species depicting the timeframe in which dosage compensation systems observed in *P. picta* and *P. parae* evolved over the ∼3.7 million year interval (denoted in green) after the common ancestor to *P. picta-P. parae* split from *P. reticulata-P*.*wingei* (∼18.4 mya) and prior to the divergence of *P. picta* and *P. parae* from each other (∼14.7 mya). The branch where sex chromosome divergence and dosage compensation evolved is indicated in green. Orange branches indicate the clade containing species where X and Y are substantially diverged and have complete dosage compensation (*P. picta*-Darolti et al 2019, *P. parae*-this study). Blue indicates species for which dosage compensation has been explicitly tested but found to be entirely lacking (Darolti et al 2019). Green star denotes the branch on which complete dosage compensation likely evolved. The phylogeny and divergence times are taken from The Fish Tree of Life (Rabosky et al. 2018).

## RESULTS

### Characterization of dosage compensation in P. parae

To test whether complete dosage compensation evolved rapidly (over ∼3.7million years) in the common ancestor of *P. picta* and *P. parae*, we performed RNA-seq on muscle tissue from males and females of *P. parae*. There are five discrete Y haplotypes in *P. parae* that segregate with the five different male morphs (immaculata, yellow melanzona, blue melanzona, red melanzona, and parae morphs). Importantly, these five *P. parae* Y haplotypes emerged after X-Y recombination was halted before the split between *P. picta* and *P. parae*, ∼ 18.4 Mya (Sandkam et al. 2020; Lindholm et al. 2004). Therefore, if complete dosage compensation evolved in the common ancestor of *P. picta* and *P. parae*, we would expect to see dosage compensation in all male morphs as well. To assess this, we tested for differences in expression from the X and Y chromosome in three of the five male morphs (yellow melanzona, blue melanzona, and parae, hereafter referred to as yellow, blue, and parae males). It is worth noting that all five Y haplotypes show similar patterns of divergence from the X (Sandkam et al. 2020), and so the three morphs we assessed here are indicative of the species as a whole.

For genes that are equally expressed from both sex chromosomes we expect to see a similar proportion of transcripts expressed from each sex chromosome. To test this, we first identified heterozygous transcripts. We found that 17% of the 38,986 autosomal transcripts exhibit heterozygous expression in both males and females, and a similar proportion (12%) of the1,349 transcripts from the sex chromosome are heterozygous in females. In contrast, only 1% of sex chromosome transcripts are heterozygous in *P. parae* males. These data suggest that widespread gene loss has occurred as a result of Y chromosome divergence in males.

We then compared the major allele ratios for heterozygous transcripts. Autosomal genes are equally expressed from both chromosomes in both sexes, and in X-linked genes in *P. parae* females (Figure 2A/B). However, in males we found significant allele specific expression (ASE) for sex-linked genes, consistent with the notion that for sex-linked genes that remain heterozygous in males, gene activity has been reduced from the Y paralog and expression is primarily produced from the X. This pattern was convergent across each male morph (Figure 2A).

**Figure 2.**
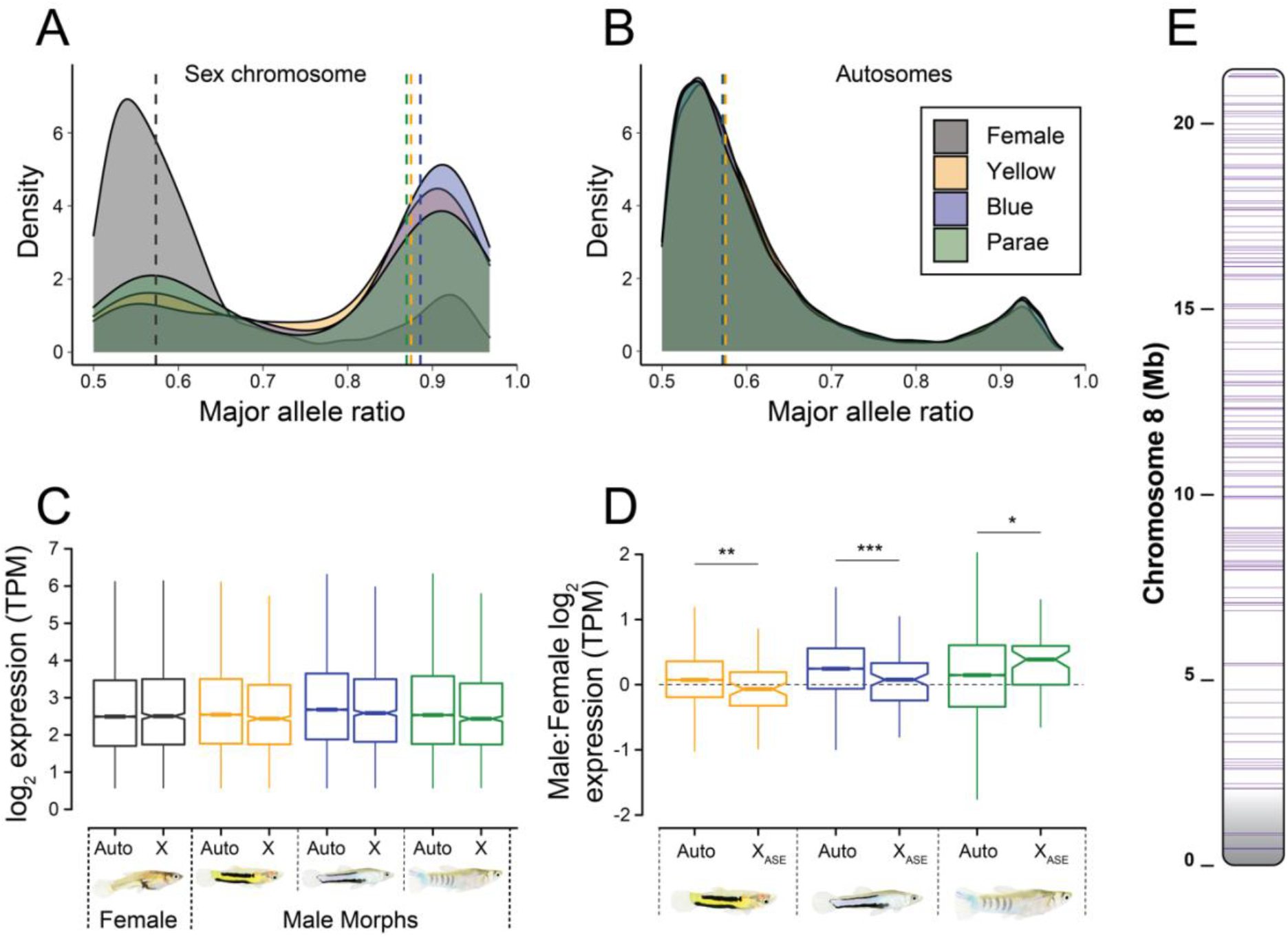
Patterns of allele specific expression (ASE) on the sex chromosome (A) and the autosomes (B) for females and each of the three male morphs of *P. parae* indicate loss of expression for genes encoded on the Y. ASE ratio of 0.5 indicates equal expression from both copies of a chromosome, while shifts toward 1 indicate expression predominantly comes from just one copy. Vertical dashed lines are median major allele ratio values. (C) Despite loss of expression from the Y, expression levels (log2 transcripts per million (TPM)) of sex chromosome genes do not differ from the autosomes for any of the male morphs. Male:female expression ratios for genes that exhibit allele specific expression are not different from male:female expression ratios of autosomal genes, demonstrating that a loss of expression from the Y chromosome in males does not result in reduced expression. The horizontal dashed line represents equal expression between males and females. Colours are consistent in all panels and denote sex and/or male morph. Grey = female, Yellow = yellow male morph, blue = blue male morph, green = parae male morph. Distribution of genes with allele specific expression (ASE) on the male X chromosome (chromosome 8). ASE genes are evenly distributed along the entire chromosome, confirming complete dosage compensation for genes on the sex chromosome. Gene locations are demarcated by purple lines. The pseudo autosomal region (PAR) is in grey.

In order to determine whether Y degeneration has been coupled with X chromosome dosage compensation, we compared average expression for all genes from the X chromosome to the autosomal gene expression levels in both sexes. We found that expression of sex chromosome genes was not different from autosomal genes in any of the male morphs (Wilcoxon rank sum yellow p-value = 0.9703, blue p-value = 0.4965, parae p-value = 0.292), or between genes on the X chromosome or autosomes in females (Wilcoxon rank sum p-value = 0.8336). Together this indicates complete dosage compensation arose before the morphs diverged and likely in the common ancestor of *P. picta* and *P. parae* (Figure 2C).

Moreover, we observe a marginal decrease in the male:female expression ratio for sex-linked genes with an ASE pattern in the yellow (p-value = 0.001) and blue (p-value = 0.0001) male morphs compared to genes on the autosomes which is consistent with the expression pattern in *P. picta* (Darolti et al., 2019). In contrast, the male:female expression ratio for sex chromosome genes with ASE in the parae male morph were significantly higher compared to the male:female ratio of autosomal genes (p-value = 0.05) (Figure 2D). These data indicate that the efficiency of the dosage compensation in *P. parae* is similar to *P. picta* and that there may be residual Y-linked expression of these genes or that dosage compensation results in a moderate overexpression of some X-linked genes in males.

In general, there are two ways in which complete dosage compensation has been observed in XY systems. In eutherian and marsupial mammals, one of the two X chromosomes is silenced in females. Although this balances sex chromosome gene expression between males and females, it does not address expression differences between X-linked and autosomes genes. In fact, X inactivation in females means that both sexes on average express X-linked genes less than the autosomal average, and only dosage sensitive genes on the X are upregulated in both sexes to counter this (Pessia et al. 2012). Alternatively, in *Drosophila* (Marín et al. 2000), and *Anolis* (Marin et al. 2017), dosage compensation is achieved by doubling the expression of genes on the X chromosome in males.

We found that expression of genes on the X chromosome is not different from expression of genes on the autosomes in females, or any of the male morphs, and that the major allele ratio for X-linked genes in females is close to 0.5 indicating roughly equal expression from both copies of the X. Furthermore, ASE genes in males are distributed along the entire X chromosome providing further support for dosage compensation of the entire chromosome (Figure 2E). Taken together, these data suggest that complete dosage compensation in *P. parae* is more similar to dosage compensation in *Drosophila* and *Anolis* where genes on the X are hyper expressed in males. This provides an excellent avenue to explore the mechanisms controlling expression across entire chromosomes.

## DISCUSSION

### Rapid evolution of dosage compensation

Although theory suggests complete dosage compensation should evolve rapidly in tandem with Y degradation (Ohno 1967), gene expression studies in non-model systems with heteromorphic sex chromosomes have demonstrated that complete dosage compensation is actually quite rare and is not a guaranteed outcome of sex chromosome evolution (Mank et al. 2011). These studies show that there are many alternatives to evolving complete dosage compensation, and that complete dosage compensation is an exceptional outcome of sex chromosome evolution. Until the recent characterization of complete dosage compensation in *P. picta*, complete dosage compensation has been observed in a limited number of lineages, all of which are relatively ancient (Marin et al. 2017; Mullon et al. 2015). The age of these systems makes it difficult to refine estimates for the speed at which complete dosage compensation can arise.

Within the family Poeciliidae the subgenus *Lebistes* is particularly well suited to address this question as it contains several species with characterized sex chromosomes including *P. reticulata, P. wingei, P. picta*, and *P. parae* (Darolti et al. 2019). There is strong evidence that all *Lebistes* share the same sex chromosome system which originated 18.48-26.08 Mya (Darolti et al. 2019; Rabosky et al. 2018). Despite sharing the same XY system, the extent of Y degradation differs dramatically, from largely intact in *P. reticulata* and *P. wingei* to highly degraded in *P. picta* and *P. parae* (Darolti et al. 2019; Sandkam et al. 2020). Without gene loss, there would be no selective pressure to evolve dosage compensation, thus it is not surprising that a dosage compensation was not found in either *P. reticulata* and *P. wingei* (Darolti et al. 2019), where there is little evidence of decreased gene activity from the Y chromosome.

In contrast, the Y chromosomes in *P. picta* and *P. parae* exhibit substantial divergence along the entire chromosome (Sandkam et al. 2020; Darolti et al. 2019). Here we present evidence for complete dosage compensation common in multiple morphs of *P. parae*. These data suggest that the dosage compensation system in *P. parae* is shared with *P. picta* and evolved over a period of less than 4 million years in their common ancestor.

In some systems, the rapid evolution of complete dosage compensation is achieved by recruiting an ancestral or pre-existing dosage compensation mechanism (Marín et al. 1996; Marin et al. 2017). In fishes, complete dosage compensation is rare, which may be the result of frequent sex chromosome turnover and a paucity of heteromorphic sex chromosomes that makes complete dosage compensation unnecessary. As such dosage compensation in fish is frequently accomplished on a gene-by-gene basis and remains overall incomplete (Shao et al. 2014; White et al. 2015; Darolti et al. 2019) with the exception of *P. picta* (Darolti et al. 2019) and *P. parae*. Further work elucidating the mechanism of X chromosome dosage compensation *P. picta* and *P. parae* will provide novel insights in the evolution of dosage compensation mechanisms.

## METHODS

### RNA isolation and sequencing

Animals used in this study were collected in Spring 2019 from natural populations in Suriname and brought to the University of British Columbia (Vancouver, BC, Canada) aquatics facility, where they were kept in 20L glass aquaria on a 12:12 day:night cycle at 26°C and 6ppt salinity (Instant Ocean Sea Salt) and fed Hikari Fancy Guppy pellets and live brine shrimp daily. Individuals were euthanized using a lethal overdose of MS-222 and muscular tail tissue was taken from the anal pore to the base of the pectoral fin. RNA was immediately isolated using RNeasy spin columns with on-column DNase treatment (Qiagen) following the manufacturer’s recommended protocol. Library preparation and 100bp paired-end sequencing was performed on an Illumina NovaSeq 6000 at McGill University and the Génome Québec Innovation Centre. Adaptor sequences were removed and reads were quality filtered and trimmed using trimmomatic (v0.36) using a sliding window of 4 bases and a minimum Phred score of 15. Reads with leading and trailing bases with a Phred score <3 were also removed. Sequencing libraries consisted of ∼88 million reads.

### Transcript Alignment and Filtering

Reads were aligned to a previously published female *P. parae* genome assembly (Sandkam et al. 2020) using the two-pass method for STAR align (v2.7.2) (Dobin et al. 2013). Alignments were sorted by coordinate and converted to BAM format using SAMtools (v1.9). To find the full list of non-redundant *P. parae* transcripts we generated GTF files for each individual using StringTie (v1.3.6) then merged all GTF files. To remove non-coding RNA (ncRNA) we first compiled a database of all ncRNAs in reference genomes of close relatives on Ensemble: *Poecilia formosa* (PoeFor_5.1.2), *Oryzias latipes* (ASM223467v1), *Gasterosteus aculeatus* (BROAD S1), and *Danio rerio* (GRCz11). We then removed all *P. parae* transcripts that BLASTed to our ncRNA database.

### Allele Specific Expression

To ensure our results are comparable to our previous results in *P. picta* we followed the same pipeline to identify allele specific expression (Darolti et al. 2019). In short, for each sex and morph we identified SNPs separately using SAMtools mpileup (v1.9) and varscan (v2.4.3) with parameters --min-coverage 2, --min-ave-qual 20, --min-freq-for-hom 0.90, and excluding triallelic SNPs. We then filtered SNPs for a minimum site coverage of 15to account for sequencing errors, and used a variable coverage filter to account for potential effects of sequencing errors due to variable coverage levels (an error rate of 1 in 100 and a maximum coverage for a given site of 100,000) (Quinn et al. 2014). We then removed SNP clusters of more than five SNPs in 100bp window to limit potential biases from read assignments to a single reference sequence (Stevenson et al. 2013).

### Expression Level

We extracted read counts using the featureCounts from the subread package (Liao et al. 2014) and the ncRNA filtered GTF file described above. Reads with low expression (less than 10% of the mean) were removed from the dataset. We then used a Wilcoxon rank sum test to compare expression levels between groups using the wilcox.test() function in R (p < 0.05).

## DATA AVAILABILITY

Illumina .fastq read files will be made publicly available on the Genbank sequence read archive upon publication of this manuscript.

## ACKNOWLEDGEMENTS

This work was supported by the Canada 150 Research Chair Program and the European Research Council (680951) to JEM. and a Banting Postdoctoral Fellowship to BAS (Natural Science and Engineering Research Council of Canada). We thank Clara Lacy for guppy illustrations and all members of the Mank lab for helpful comments and suggestions.

